# Chronic alcohol consumption dysregulates innate immune response to SARS-CoV-2 in the lung

**DOI:** 10.1101/2023.05.02.539139

**Authors:** Sloan A. Lewis, Isaac R. Cinco, Brianna M. Doratt, Madison B. Blanton, Cherise Hoagland, Michael Davies, Kathleen A. Grant, Ilhem Messaoudi

**Author notes:** Corresponding Author: Ilhem Messaoudi Microbiology, Immunology and Molecular Genetics College of Medicine University of Kentucky, Lexington, KY. **Author contributions** S.A.L., K.A.G., and I.M. conceived and designed the experiments. S.A.L., C.H., N.N, M.D. and B.D., performed the experiments. S.A.L., I.C., and M.B. analyzed the data. M.B., I.C., and I.M. wrote the paper. All authors have read and approved the final draft of the manuscript.^1^.

## Abstract

Alcohol consumption is widespread with over half of the individuals over 18 years of age in the U.S. reporting alcohol use in the last 30 days. Moreover, 9 million Americans engaged in binge or chronic heavy drinking (CHD) in 2019. CHD negatively impacts pathogen clearance and tissue repair, including in the respiratory tract, thereby increasing susceptibility to infection. Although, it has been hypothesized that chronic alcohol consumption negatively impacts COVID-19 outcomes; the interplay between chronic alcohol use and SARS-CoV-2 infection outcomes has yet to be elucidated. Therefore, in this study we investigated the impact of chronic alcohol consumption on SARS-CoV-2 anti-viral responses in bronchoalveolar lavage cell samples from humans with alcohol use disorder and rhesus macaques that engaged in chronic drinking. Our data show that in both humans and macaques, the induction of key antiviral cytokines and growth factors was decreased with chronic ethanol consumption. Moreover, in macaques fewer differentially expressed genes mapped to Gene Ontology terms associated with antiviral immunity following 6 month of ethanol consumption while TLR signaling pathways were upregulated. These data are indicative of aberrant inflammation and reduced antiviral responses in the lung with chronic alcohol drinking.

## Introduction

Alcohol consumption is widespread in the United States, with 55% of the American population over 18 years of age reporting alcohol use within the last 30 days. Alarmingly, 25% and 6.3% of adults over 18 years of age are classified as binge or heavy drinkers, respectively (National Survey on Drug Use and Health 2020). Both binge and heavy drinking can contribute to acquiring an Alcohol Use Disorder (AUD) diagnosis, defined as the inability to control or cease alcohol use despite experiencing negative social, occupational, or health-related consequences (1). In 2019, 9 million men and 5.5 million women were diagnosed with AUD (2). Excessive alcohol consumption impairs lung (3), liver (4), pancreas (5), spleen (6), and heart (7, 8) function. Alcohol consumption is also associated with increased incidence of cancers (9–11). Moreover, excessive alcohol consumption leads to an increased rate (12) and length of hospitalizations (13) including to the intensive care unit (14). Individuals with AUD are also at increased risk of nosocomial infections following trauma (15) and wound-related infections (16). Overall, excessive alcohol use in the United States is associated with a $249 billion loss to the American economy (17) and $28 billion in healthcare-related costs (17, 18). Moreover, it is also associated with more than 260 alcohol-related deaths per day, making AUD the third leading cause of preventable deaths in the United States (19).

Data from clinical and experimental studies indicate that chronic excessive alcohol consumption disrupts cellular mechanisms responsible for pathogen clearance and tissue repair (20, 21), leading to increased susceptibility to infection and delayed wound healing (22, 23). Specifically, the incidence of K. pneumonia, S. pneumonia, Mycobacterium tuberculosis infection, hepatitis C virus, and respiratory syncytia virus (RSV) are increased with chronic heavy drinking (CHD) (23–25). Animal model studies have recapitulated the increased severity of bacterial and viral infections with CHD further highlighting the immunological basis of these adverse outcomes (26, 27).

Underlying mechanisms driving poor infectious disease outcomes are the focus of several studies that have reported hyper-inflammatory responses following LPS stimulation by circulating and tissue-resident myeloid cells from humans with or animal models of CHD (3, 28). In contrast, stimulation with pathogens decreased the production of key immune mediators (3, 29, 30). A dysregulation of phagocytic capacity within myeloid populations has also been observed in conjunction with CHD (31, 32). Additionally, in-vitro ethanol treatment of monocytes resulted in lower expression levels of MHC-II molecules (33), suggesting that CHD can lead to reduced antigen presentation capacity. Similarly, excessive alcohol consumption interferes with the phagocytic capacity of alveolar macrophages (AM) and their ability to generate anti-viral and anti-bacterial responses (34–36). Alcohol exposure in rats dampens the expression of GM-CSF receptors required for the differentiation of monocytes into macrophages upon infiltrating the lungs (34, 37). In addition to impairing immune responses, CHD decreases barrier function in the respiratory tract, increasing susceptibility to respiratory infections (38). Individuals with CHD are at increased risk of developing acute respiratory distress syndrome (ARDS) (39), leading to organ failure, sepsis, or death (40).

The shelter-in-place orders associated with the COVID-19 pandemic led to increased alcohol consumption in the United States (41). Given the impact of chronic ethanol consumption on the respiratory system, it has been hypothesized that increased ethanol consumption could be one of the drivers for COVID-19 severity (42). However, the interplay between chronic ethanol consumption and the anti-viral response to SARS-CoV-2 remains poorly understood. Therefore, in this study, bronchial alveolar lavage (BAL) cell samples obtained from rhesus macaques before and after 6-months of voluntary ethanol self-administration, as well as those obtained from individuals with and without AUD were infected ex vivo with SARS-CoV-2. Response to infection was monitored by measuring immune mediator release and transcriptional changes using single cell RNA sequencing (scRNA-Seq).

## Materials and Methods

See the data supplement for detailed methods

### Sample collection

Bronchoalveolar lavage (BAL) samples were collected from 5 male and 6 female rhesus macaques (6 low drinkers, 4 heavy drinker, and 1 very heavy drinker) at baseline (before induction) and after 6 months of chronic consumption (**Table 1**) (Cohort 18 on matrr.com) (43, 44).

**Table 1:** Demographics of the NHP cohort utilized in this study.

| ID | Sex | Drinking<br>Phenotype | Mean daily<br>ethanol intake<br>(g/kg/day) | Average Blood<br>Ethanol<br>Content (Avg<br>mg%) |
| --- | --- | --- | --- | --- |
| 10331 | F | LD | 0.5 | 0 |
| 10332 | F | LD | 1.23 | 3.15 |
| 10334 | F | LD | 1.78 | 24.32 |
| 10339 | F | LD | 1.74 | 30.42 |
| 10340 | F | VHD | 3.53 | 40.24 |
| 10333 | F | HD | 3.04 | 46.01 |
| 10341 | M | LD | 1.94 | 32.64 |
| 10335 | M | LD | 1.32 | 16.78 |
| 10337 | M | VHD | 4.02 | 158.62 |
| 10338 | M | VHD | 5.93 | 209.11 |
| 10342 | M | VHD | 4.04 | 161.72 |

Human BAL samples and informed written consent were obtained, after IRB approval, from the University of Colorado School of Medicine’s Colorado-Pulmonary Alcohol Research Collaborative (CoPARC) (https://medschool.cuanschutz.edu/coparc) (**Table 2**).

**Table 2:** Demographic information from the human BAL samples utilized in this study.

|  | Smoking<br>n=3 | Smoking and Alcohol Constumption<br>n=3 |
| --- | --- | --- |
| Age (years) | 54.7 ± 6.1 | 58.0 ± 2.6 |
| Sex | Males: 3<br>Females: 0 | Males: 3<br>Females: 0 |
| <u>Race</u> |  |  |
| African American | 1 | 1 |
| White | 1 | 1 |
| Hispanic | 1 | 1 |

### Bronchoalveolar Lavage (BAL) Cell Isolation

BAL samples were centrifuged at 2000 rpm for 10 minutes. Pellets were resuspended in 10% DMSO/FBS solution, stored in a Mr. Frosty Freezing container (Thermo Fisher Scientific) at -80C for 24 hours, and then transferred to a cryogenic unit for long-term storage.

### Flow Cytometry

BAL cells were thawed, stained with the specified antibodies, acquired using an Attune NxT Flow Cytometer, and analyzed using the FlowJo software.

### SARS-CoV-2 Infection

1x10^6^ live BAL cells were infected with SARS-CoV-2 (Washington isolate) at an MOI of 1 or left untreated for 24 hours at 37°C, 5% CO_2_. At the end of the infection, supernatants were stored at -80C for Luminex analysis while cells were processed for single cell RNA sequencing.

### Luminex

Immune mediators were measured in NHP samples via an 36-plex NHP XL Cytokine Premixed Kit (R&D) and in human samples via a customized human 29-plex (R&D). The samples were analyzed on a MAGPIX instrument (Luminex).

### Single Cell RNA library preparation

Live BAL cells were sorted and labeled with cell multiplexing oligos per manufacturer instructions (10X Genomics) then loaded into a 10x Genomics Chromium Controller. The libraries were prepared using the v3.1 chemistry Single Cell 3’Feature Barcoding Library Kit (10x Genomics) according to the manufacturer’s instructions.

### Single cells RNA Sequencing Analysis

Libraries were sequenced using a NovaSeq6000. The reads were aligned and quantified against the Mmul_8 rhesus macaque reference genome or the human genome GRCh38. Downstream analysis was performed using Seurat *multi-*function (version 4.1.1). Samples were de-multiplexed (45), normalized, and integrated. A PCA was generated to determine the dimensionality of the data set and a UMAP was generated and cell clustering was performed. Differential gene expression analysis and functional enrichment were performed using MAST and Metascape, respectively (46).

### Data availability

The datasets utilized in this article are available on NCBI’s Sequence Read Archive PRJNA930878 (NHP) and PRJNA930381 (Human).

### Statistical Analysis

Statistical analyses were carried out using GraphPad Prism 9.

## Results

### Chronic heavy drinking skews the lung environment towards a heightened inflammatory state

To understand the impact of chronic ethanol consumption on the immunological landscape of the lung, BAL samples were collected from male and female rhesus macaques before and after 6 months of daily voluntary ethanol self-administration (**Table 1**) and subjected to flow cytometry (**Figure 1A, Supp Figure 1 A, B)**. This analysis revealed that at baseline the relative abundance of CD8 and CD4 T cells was significantly higher in females compared to males (**Supp Figure 1C**). In contrast, the frequency of B cells was higher in males than females at baseline (**Supp Figure 1D**). Following 6 months of ethanol self-administration, the frequency of CD8 T cells was significantly reduced in females while that of natural killer (NK) cells was significantly reduced in males (**Supp. Figure 1C, E**). Conversely, the relative abundance of alveolar macrophages (AM) was increased in both males and females (**Supp. Fig 1F**). This increase was primarily mediated by an expansion of activated AM (CD163+), and a decrease in interstitial macrophages (IM) (**Supp. Fig 1F, G**). The frequency of dendritic cells (DC) also significantly increased after 6 months of CHD in males, whereas the opposite trend was observed in females (**Supp. Fig 1H**). The frequency of infiltrating monocytes was greater in females than males at baseline but decreased with drinking (**Supp. Fig 1I**). Nevertheless, ethanol exposure led to an expansion of nonclassical (activated) monocytes and a decrease in classical monocytes in both males and females (**Supp. Fig 1I**).

**Figure 1:**
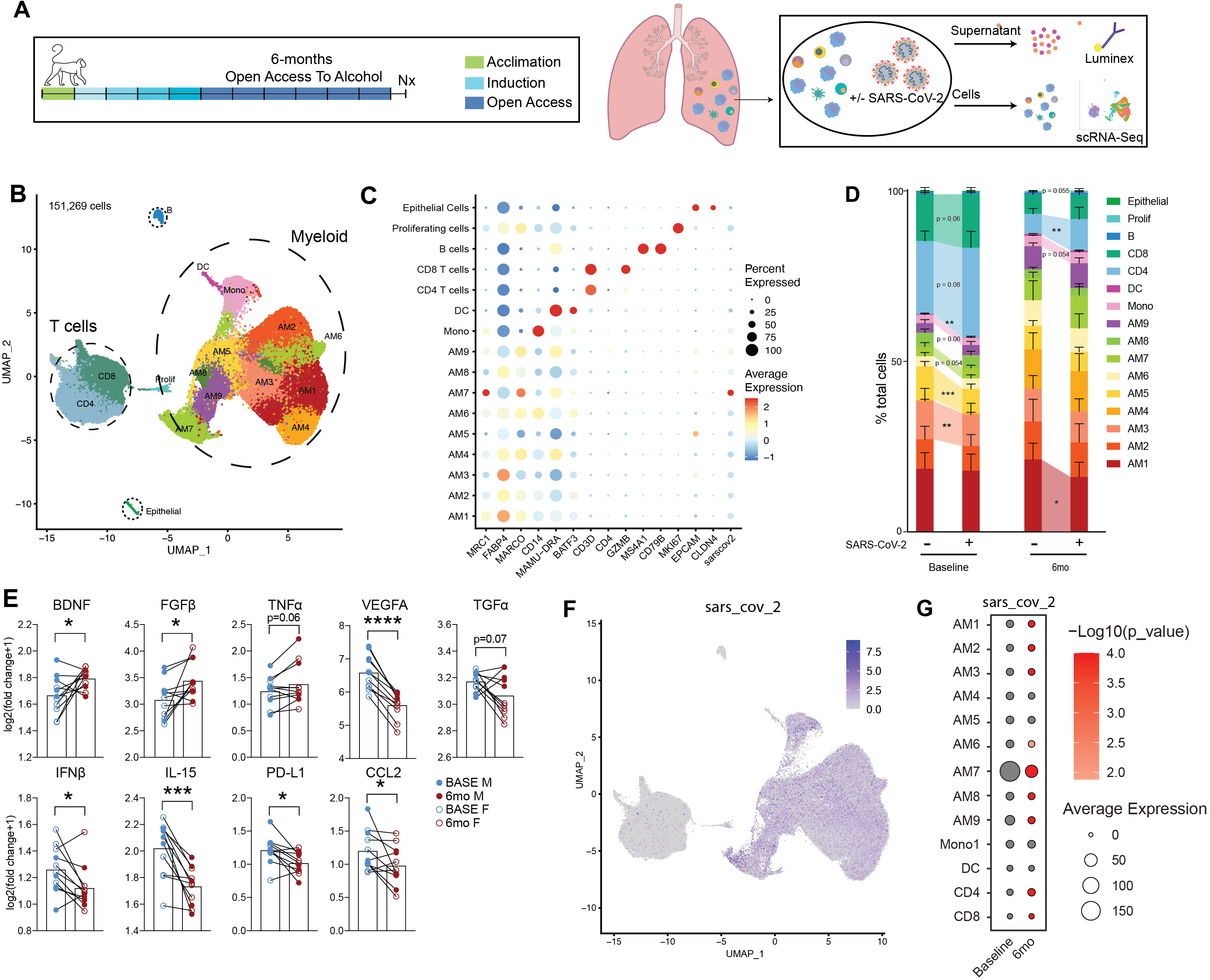
Response to SARS-CoV-2 infection is mitigated in rhesus macaques after chronic alcohol consumption. **A)** BALs were collected from rhesus macaqus before and after 6 months of drinking to assess the impact of alcohol consumption on SARS-CoV-2 infection. B) Uniform manifold approximation and projection (UMAP) representation of 151, 269 immune cells with and without infection and 6 months drinking. Sixteen unique clusters were determined. C) Bubble plot of key cluster marker genes. Percent of cells that express each transcript is represented by the size of the bubble while the average expression of those cells is denoted by color, ranging from blue (low) to red (high). **D)** Relative frequency of each cluster before and after infection. Samples within time points were compared in a pairwise fashion. **E)** Scatter bar plots comparing the Log2 fold change in response to SARS-CoV-2 infection of select cytokine and chemokine concentrations within BAL cell culture supernatants at baseline and after 6 months of open access to alcohol in male and female subjects (* p ≤ 0.05, ** p ≤ 0.01, *** p ≤ 0.001, **** p ≤ 0.0001). **F)** Feature plot of SARS-CoV-2 expression within the UMAP. **G)** Bubbleplot of SARS-CoV-2 expression within each cluster stratified by timepoint. Average expression of the SARS-CoV-2 transcript is represented by bubble size while the significance of baseline to 6 months comparison is denoted by color.

We next investigated the impact of chronic ethanol consumption on the transcriptional landscape of immune cells using scRNA-Seq (**Figure 1A**). A total of 16 unique clusters were identified, including 9 alveolar macrophages (AM) subsets (defined by markers *MRC1, FABP4, MARCO*), 1 infiltrating monocyte group (*CD14*), 1 conventional DC subset (*MAMU-DRA, BATF3*), CD4 and CD8 T cells (*CD3D, CD4, GZMB*), B cells (*MS4A1, CD79B*), proliferating cells (*MKI67*), and epithelial cells (*EPCAM, CLDN4*) populations (**Figure 1B, C**). Each cluster was constituted of cells from all subjects, regardless of the time point of collection and SARS-CoV-2 infection (**Supp. Fig 2A**). The 9 AM clusters were distinct based on several inflammatory and regulatory markers (*FABP4, STAT1, CD9.1, MMP10, ISG20, IL1B, CD14, MARCO, LCN2, HYOU1, LYZ, MAMU-DRA*) (**Supp. Fig 2B**).

In line with the data from flow cytometry, the relative abundance of T cells decreased while that of myeloid cells increased following 6 months of ethanol consumption (**Figure 1D**). Ethanol consumption also resulted in transcriptional shifts in the immune landscape of the lung. Specifically, within myeloid populations (AM, infiltrating monocytes), scores of modules associated with migration were decreased while those associated with hypoxia (HIF1α signaling) and other inflammatory pathways (TLR, viral/bacterial responses) were increased **(Supp. Fig 3A)**. Similarly, genes differentially expressed (DEG) with alcohol consumption in myeloid subsets mapped to GO terms such as response to oxygen levels (*HlF-1α*), innate immune responses (*CD14*), and positive regulation of cell migration (*IL-1α, IL-1β, MMP14, CD74, MIF*) **(Supp. Fig 3B, C)**. Within CD4 and CD8 T cell subsets, scores of modules associated with Th2 and viral/bacterial response signaling pathways were increased with chronic ethanol consumption **(Supp. Fig 3A)**. DEGs in lymphoid populations map to GO terms such as cell killing (*GZMB*), lymphocyte-mediated immunity (IL-2R*β*, *CD40LG*), in addition to response to hypoxia **(Supp. Fig 3B, C)**. These data indicate that chronic ethanol consumption alters the frequency and transcriptional profile of BAL-resident immune cells.

### Chronic heavy drinking alters the immune mediator response to SARS-CoV-2 infection and distribution of cell subsets

We next assessed the impact of ethanol consumption on immune mediator production in response to SARS-CoV-2 infection (**Figure 1A**). The increase in the production of immune mediators over no infection condition was comparable between males and females (**Supp Table 1**) so the data were combined. SARS-CoV-2 infection induced a robust response demonstrated by increased induction of several cytokines, chemokines, and growth factors in culture supernatants at both timepoints relative to non-infection conditions (**Figure 1E, Supp. Table 1)**. Chronic ethanol consumption resulted in an increase in the induction of growth factors BDNF and FGFβ and a modest increase in the induction of inflammatory cytokine TNFα. In contrast, induction of growth factors VEGFA and TGFα, inflammatory cytokines IFNβ and IL-15, T cell activation marker PD-L1, and chemokine CCL2 were significantly decreased after 6 months of drinking (**Figure 1E**). These results demonstrate that chronic ethanol consumption alters the secrotome in response to viral infection, suggestive of a dysregulated SARS-CoV-2 responses.

We then leveraged scRNA Seq to investigate the impact of chronic ethanol consumption on immune subset distribution and SARS-CoV-2 detection. A pairwise comparison of the relative abundance of clusters after infection showed a significantly reduced frequency of the AM3, AM5, and DC clusters and a notable decrease in the AM7, Mono, CD4, and CD8 clusters at baseline (**Figure 1D**). After 6 months of ethanol consumption, significant decrease in frequency was evident in AM1 and CD4 clusters, with a notable decrease in the Mono cluster and an increase in the Epithelial cluster was observed in response to SARS-CoV-2 infection (**Figure 1D**). Additionally, this analysis revealed that myeloid clusters contained a majority of the SARS-CoV-2 transcript, with the AM7 cluster exhibiting the greatest expression (**Figure 1F,G**). Interestingly, the average expression of SARS-CoV-2 transcripts was significantly decreased across all AM clusters, except AM4 and AM5, after 6 months of ethanol consumption (**Figure 1G**). In contrast, levels of viral transcripts were increased in the CD4 and CD8 T cell subset after 6 months of chronic ethanol consumption (**Figure 1G**).

### Chronic ethanol consumption alters transcriptional response to SARS-CoV-2

Because SARS-CoV-2 was predominately detected in myeloid cells, we focused our analysis on these clusters. To interrogate the impact of ethanol consumption on the anti-SARS-CoV-2 response, we measured module scores associated with cytokine/chemokine signaling, viral/bacterial response, and toll-like receptor (TLR) signaling (**Figure 2A, Supp Table 2**). All three module scores increased after SARS-CoV-2 infection at both time points within several myeloid clusters (**Figure 2A**). After infection, there was a significant increase in module scores of cytokine/chemokine signaling and anti-viral/bacterial responses within more myeloid clusters at baseline, whereas scores associated with TLR signaling were increased within a greater number of clusters after ethanol consumption (**Figure 2A**). Futhermore, module scores for cytokine/chemokine signaling were significantly lower in AM3, AM4, and Mono at 6mo when compared to baseline after infection. These observations are in line with the decreased induction of IL-15, VEGF, IFNβ and increased induction of TNFa observed in the Luminex analysis (**Figure 1E**). All three module scores increased in DCs after infection in the samples obtained at baseline whereas a significant increase in the expression of genes only within the anti-viral/bacterial module were noted after 6 onths of ethanol consumption (**Figure 2A**).

**Figure 2:**
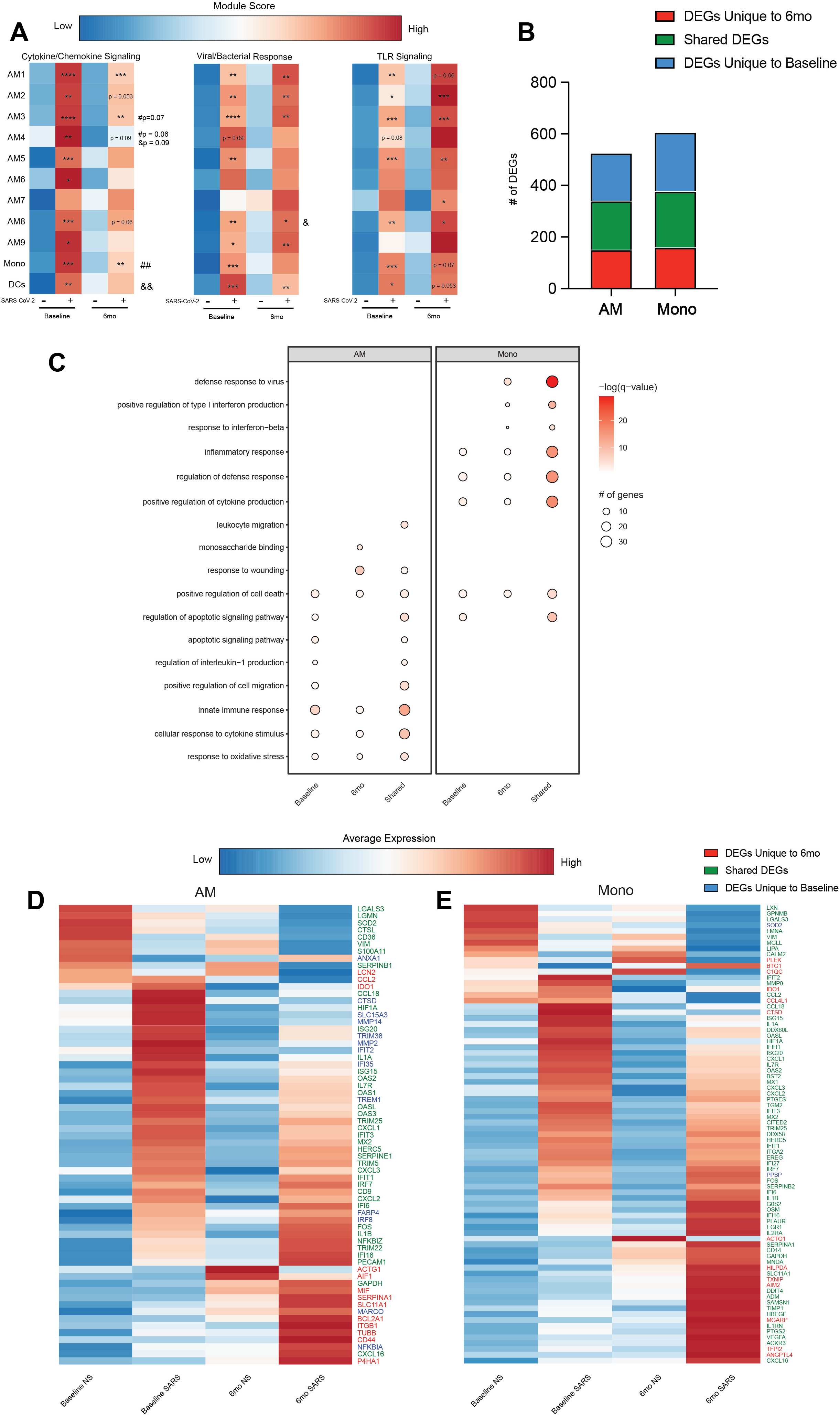
CHD predominately affects how myeloid cells respond to SARS-CoV-2. **A)** Heatmap comparing module scores for cytokine/chemokine signaling, viral/bacterial responses, and TLR signaling between samples based on condition and timepoint. The average module score for each cluster ranges from low (blue) to high (red). Significance displayed is relative to non-infected conditions within timepoints, * p ≤ 0.05, ** ≤ 0.01, *** p ≤ 0.001, **** p ≤ 0.0001. # denotes significance when comparing infected conditions and & denotes significance when comparing non infected conditions (#/& p ≤ 0.05, && ≤ 0.01). **B)** Stacked bar graph showing the number of unique and shared DEGs for myeloid clusters. **C)** Bubble plot of significant and notable gene ontology (GO) terms from functional enrichment of myeloid clusters. **D)** Heatmap of the average expression of select DEGs in AM clusters across conditions and timepoint. The average expression for each transcript ranges from low (blue) to high (red). **E)** Heatmap of the average expression of select DEGs in mono clusters across conditions and timepoint. The average expression for each transcript ranges from low (blue) to high (red).

We next performed differential gene expression analysis comparing infected relative to noninfected cells within each time point. For this analysis, the AM clusters were collapsed. SARS-CoV-2 infection was found to be associated with a robust transcriptional response both at baseline and after 6mo of chronic drinking (**Figure 2B**). While a significant overlap between was noted between the two time points, many DEGs unique to baseline or post-ethanol consumption conditions were also identified (**Figure 2B**). DEGs unique to baseline enriched to processes associated with innate immunity and apoptosis (e.g. “regulation of IL1 production” and “apoptotic signaling pathway”) (**Figure 2C**). Some of the notable unique DEG include cathepsin D (*CTSD*), metalloproteases (*MMP2* and *MMP4*) as well viral restriction factors (*TRIM36*, *IFI35*) and regulatory molecules (*TREM1*) (**Figure 2D**). On the other hand, DEGs unique to post ethanol consumption enriched to gene ontology (GO) terms “monosaccharide binding” and “response to wounding” including *MIF*, *SERPINA1*, *LCN2*, *CCL2*, and *IDO1* (**Figure 2C,D**). Within the infiltrating monocytes, a greater number of genes enriched to GO terms “defense response to virus” and “positive regulation of type I IFN” (**Figure 2C**). DEG unique to the 6mo time point include *TXNIP*, *IDO1*, *CC4L1*, and *C1Q* in line with heightened TLR signaling pathways (**Figure 2E**).

Although T cells were not the primary target of SARS-CoV-2 (**Fig 1F**), transcriptional responses to infection were noted **(Supp. Fig 2C, D)**. DEGs that play a role in type I and II interferon and anti-viral pathways were more evident at baseline. Those detected only after chronic drinking enriched primarily to apoptotic pathways **(Sup Fig 2C, D)**. Overall, these findings indicate that ethanol consumption skews the transcriptional profiles of alveolar cells towards a hyperinflammatory state while antiviral responses are dampened.

### Dysregulation of anti-SARS-CoV-2 response by AUD in humans

To place the observations obtained from the NHP model after 6 months of drinking in the context of long-term chronic heavy drinking in humans, BAL samples were obtained from individuals with AUD and smoking or smoking only (n=3/group) (**Table 2**). Flow cytometry revealed a significant increase in the B cell population within the control group compared to the AUD group (**Table 3**). Transcriptional responses to ethanol consumption were assessed by scRNA-Seq. Using canonical gene markers, we identified 6 AM clusters (*MARCO, HLA-DMA, C1QA, GCGR3A, FABP4, OASL*) with one of the clusters being a Proliferating AM cluster (*MKI67*), 2 infiltrating monocyte clusters (*CCL2, CXCL8, CD14,*), 1 DC cluster (*CCR7, LAMP5*), and 1 CD8 T cell cluster (*CD3D, CD8A, CD8B, GZMA*) (**Figures 3A, B**). The clusters were distributed among all samples and timepoints (**Supp. Fig 4A**). The 5 non-proliferating AM clusters were found to be distinct based on unique expression patterns of *CXCL15, IL1B, FOS, CD14, SLC3A2, ARK1C3, FN1, FBP1, LIPA, CAQB, CTSD, HLA-DQB1, CXCL8, ATF3, OASL* (Supp. Fig 4B).

**Table 3:** Frequencies of immune cell populations determined by flow cytometery

| Cell type | Control (%) |  |  | AUD (%) |  |  | Derived |
| --- | --- | --- | --- | --- | --- | --- | --- |
|  | 1 | 2 | 3 | 1 | 2 | 3 |  |
| AM | 61.7 | 74.1 | 80.9 | 72.6 | 86.1 | 53.6 | % of Live Cells |
| AM 163- | 90.5 | 90.8 | 95.3 | 85.7 | 89 | 90.2 | % of Live AM |
| AM 163+ | 6.55 | 6.91 | 3.02 | 12.3 | 9.1 | 5.67 | % of Live AM |
| IM | 5.87 | 5.21 | 5.03 | 7.76 | 4.38 | 12.6 | % of Live Cells |
| DC | 1.88 | 2.26 | 0.37 | 3.15 | 0.6 | 1.33 | % of Live Cells |
| Monocytes | 1.18 | 3.09 | 0.79 | 3.74 | 0.51 | 1.71 | % of Live Cells |
| Classical Monocytes | 97.6 | 98.3 | 97 | 98.9 | 95.4 | 94.5 | % of Live Monocytes |
| Non-classical Monocytes | 2.41 | 1.67 | 2.95 | 1.09 | 4.6 | 5.54 | % of Live Monocytes |
| CD4+ | 15.4 | 18.4 | 5.99 | 17.1 | 30.9 | 8.96 | % of Live Cells |
| CD8+ | 27.3 | 17.2 | 5.55 | 27.6 | 4.8 | 9.47 | % of Live Cells |
| NK | 1.48 | 2.52 | 2.69 | 2.4 | 1.67 | 3.18 | % of Live Cells |
| B Cells* | 0.74 | 1.01 | 0.67 | 0.15 | 0.38 | 0.46 | % of Live Cells |

\* significant difference in cell populations when comparing the control group to the AUD group.

Similar to NHP, ethanol consumption resulted in transcriptional alteration of immune cell populations within the BAL. There was an increase in scores of modules associated with migration, oxidative stress, hypoxia, and signaling pathways (TLR, cytokine/chemokine, NFKB, HIF-1α) within myeloid cell populations (AM and monocytes) **(Supp. Fig 5A)**. Furthermore, DEGs in myeloid populations after ethanol consumption mapped to GO terms such as inflammatory response (*CXCL1, CXCL3, FN1, LYZ)* and positive regulation of cytokine production (*CD36, CD84, IL-1α)* **(Supp. Fig 5B,C)**. DEGs such as *GNLY*, *CCL4*, and *GZMB* were detected within lymphoid subsets (CD8) **(Supp. Figure 5C)**.

Following SARS-CoV-2 infection, AUD was associated with modest increases in the secretion of cytokines, chemokines, and growth factors (PDGF-BB, IL-7, VEGF, IL-18, IL-15, CXCL10, CCL11) whereas infection of BAL cells from non-AUD individuals did not lead to a significant production of immune mediators (**Figure 3C, Supp Table 1**). Infection with SARS-CoV-2 did not lead to significant alterations in cluster frequency other than a modest increase in AM1 in the control group (**Figure 3D**).

**Figure 3:**
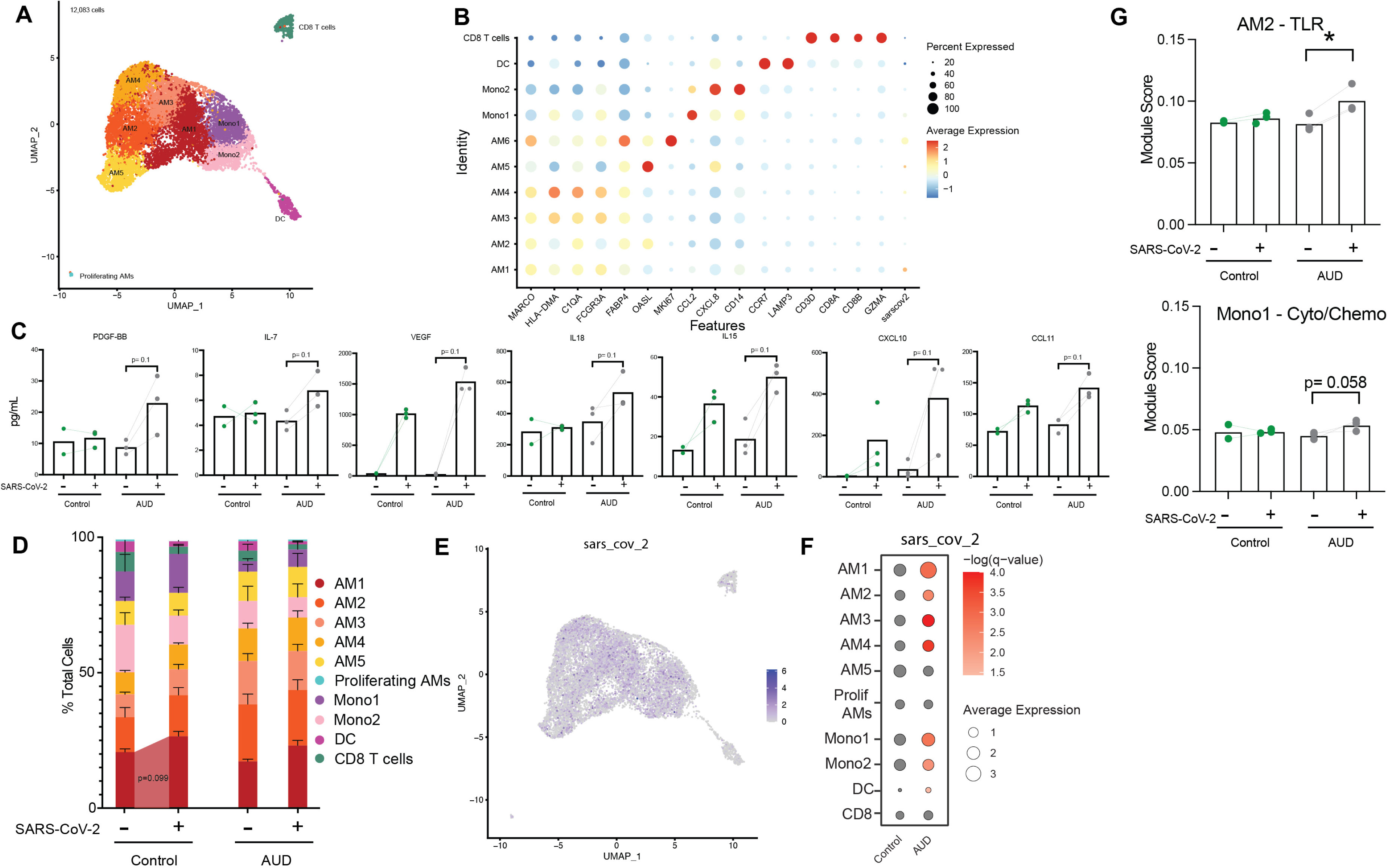
Alcohol consumption in humans dysregulates immune responses to SARS-CoV-2. **A)** UMAP visualization of 12,083 cells from both control and alcohol cohorts prior to and after SARS-CoV-2 stimulation**. B)** Bubble plot of canonical marker genes used to identify cell clusters. Percent of cells that express each transcript is represented by the size of the bubble while the average expression of those cells is denoted by color, ranging from blue (low) to red (high). **C)** Barplot of immune mediator production stratified by condition and timepoint (trending p ≤ 0.1) (EtOH non-infected, Control SARS, and EtOH SARS n=3; Control non-infected n=2). **D)** Cluster frequency stratified by condition and timepoint. **E)** Average expression of SARS-CoV-2 in the UMAP and compared between control and AUD groups. **F)** Bubbleplot of SARS-CoV-2 expression within each cluster stratified by timepoint. Average expression of the SARS-CoV-2 transcript is represented by bubble size while the significance of expression when comparing baseline to 6 months is denoted by color. **G)** Comparison of module scores of genes mapped to TLR signaling, cytokine/chemokine signaling, and viral/bacterial response, stratified by stimulation condition and collection timepoint. (* p ≤ 0.05, trending p ≤ 0.1).

As described for the macaque samples, SARS-CoV-2 transcripts were primarily detected in AM and infiltrating monocyte populations (**Figure 3E, F**). In all AM clusters, except for AM5 and proliferating AM, the expression of SARS-CoV-2 transcripts was higher in the AUD group compared to controls with viral transcripts more abundant in Mono1, Mono2, and DC (**Figure 3E, F**). Module scores associated with TLR and cytokine/chemokine signaling were increased in the SARS-CoV-2 infected AUD samples of AM2 and Mono2 cells respectively (**Figure 3G**). Overall, DEGs in the AUD group played critical roles in anti-viral, inflammatory, and oxidative stress responses (**Supp. Fig 4C**). Interestingly, several genes had different expression patterns with infection in AUD samples including ISG (*ISG20, IFI6*), cytokines (*IL6, TNF),* and chemokines (*CXCL2, CXCL3, CCL22, CXCL10*) (**Supp. Figure 4D**).

## DISCUSSION

Chronic heavy drinking has been associated with increased incidence and severity of respiratory infections (47–49). Shelter-in-place orders during the COVID-19 pandemic have led to increased alcohol consumption in the United States (41). It has been hypothesized that chronic alcohol consumption negatively impacts COVID-19 outcomes (50). However, this question has yet to be investigated in greater detail. Thus, in this study, we investigated the cellular and transcriptional impact of chronic alcohol consumption on responses to SARS-CoV-2 infection using BAL samples from female and male rhesus macaques that self-administered ethanol for six months (6, 51–54) and human male subjects with AUD.

In macaques, most immune cell populations decreased in relative frequency following six months of ethanol consumption, with the exception of AM. The increase in the relative frequency of AM was more pronounced in male macaques, which is in accordance with data from rodent models (55). Moreover, ethanol consumption led to a shift toward inflammatory AM and non-classical monocytes in line with heightened inflammation (3, 28, 51, 56). We also report that chronic ethanol consumption skews the immune landscape towards inflammatory responses, as indicated by increased expression of genes within the HIF1α as well as TLR signaling pathways.

SARS-CoV-2 transcripts were primarily detected within myeloid subsets in BAL. These observations align with earlier studies that have reported the detection of SARS-CoV-2 viral RNA or transcripts in BAL (57–59) as well as scRNA-seq analysis of whole lung tissue that reported SARS-CoV-2 viral reads within macrophage, endothelial, DC, and monocyte populations (59). Studies have also shown that AM can express ACE2 (60, 61) and TMPRSS (61), both required for SARS-CoV-2 infection. However, it remains unclear whether monocytes/macrophages can support SARS-CoV-2 replication or whether the presence of the SARS-CoV-2 transcripts is simply a consequence of phagocytosis of dying infected cells or viral particles (62).

In line with the detection of SARS-CoV-2 transcripts, the most significant transcriptional differences after infection were detected within myeloid subsets. Chronic ethanol consumption led to greater induction of TLR signaling pathways in line with previous studies that showed chronic inflammation increases oxidative stress and activates TLR signaling cascades (63, 64). In contrast, after 6 months of chronic drinking, the induction of genes important for anti-viral responses was reduced across multiple subsets compared to baseline. Reduced ISG expression could be a result of reduced production of IFN*β*. These data are similar to those from previous studies using the same rhesus macaque model of self-administration, which reported heightened pro-inflammatory responses and dampened expression of ISG in AM subsets in response to RSV infection (3). Decreased production of immune mediators involved in tissue repair mechanisms, such as VEGF, suggests a decreased tissue repair capacity (65). Reduced antiviral responses and tissue repair capacity may explain the exacerbated SARS-CoV-2 disease severity documented among AUD patients (66).

Although T cells harbored very few SARS-CoV-2 transcripts, the expression of genes important for chemotaxis, anti-viral responses, cytokine signaling, and apoptosis were altered. These transcriptional responses are likely secondary to the inflammatory responses generated by the myeloid cells after CHD and align with previous reports of impaired T cell function after alcohol exposure (67–69). The lack of induction of genes mapping to leukocyte migration could be explained by the decrease in T-cell chemoattractant IL-15 following CHD (70).

Myeloid cells were also the predominant cell type to harbor SARS-CoV-2 transcripts in BAL samples from humans with AUD. However, inflammatory mediators were produced at greater levels in samples from individuals with AUD compared to the control smoking group only. This differs from the data obtained with NHP control BAL samples which were obtained before beginning of ethanol self-administration, suggesting that smoking may abrogate inflammatory responses. Indeed, previous work has shown that tobacco smoking reduces both gene expression and production of proinflammatory cytokines (*TNFα, IL-1β,* and *IL-16*) in alveolar macrophages stimulated with *TLR2* and *TLR4* agonists (71). Additional studies utilizing electrochemiluminescence techniques to measure immune mediator production indicated that CHD, without concomitant smoking, produced the most robust inflammatory cytokine response (*IFN-γ, IL-1β,* and *IL-6*) and smoking with AUD resulted in decreased AM immune mediator responses to bacterial stimuli (72).

In summary, we show that CHD results in the dysregulation of SARS-CoV-2 response pathways. The innate branch of the immune system has proven to be a key component in modulating both SARS-CoV-2 disease severity and clinical outcomes (73). CHD reprograms innate immune cells, skewing their ability to respond to pathogens appropriately and efficiently by decreasing antiviral cytokines and transcriptionally altering the viral recognition and response pathways. Such reprogramming can induce aberrant inflammation and reduced antimicrobial responses (53). Our study supports this conclusion with the demonstration that impaired response to pathogens involves CHD-induced transcriptional changes that then increase both susceptibility and severity of SARS-CoV-2 infection.

## Acknowledgments

We could like to thank Dr. Delphine Malherbe for critical reading of the manuscript, all the members of Dr. Kathy Grant for expert aniamal care, the Division of Comparative Medicine at the Oregon National Primate Center for sample collection, and Dr. Ellen Burnham’s team for providing human BAL samples.

## Competing interests

There are no competing interests to report.

**Supp Table 1: Raw Luminex Data**: https://data.mendeley.com/datasets/f44h3v2fn8

**Supp Table 2: Gene lists for Module Scores**: https://data.mendeley.com/datasets/729m9vxf4v

---

**Supplemental Figure 1:**
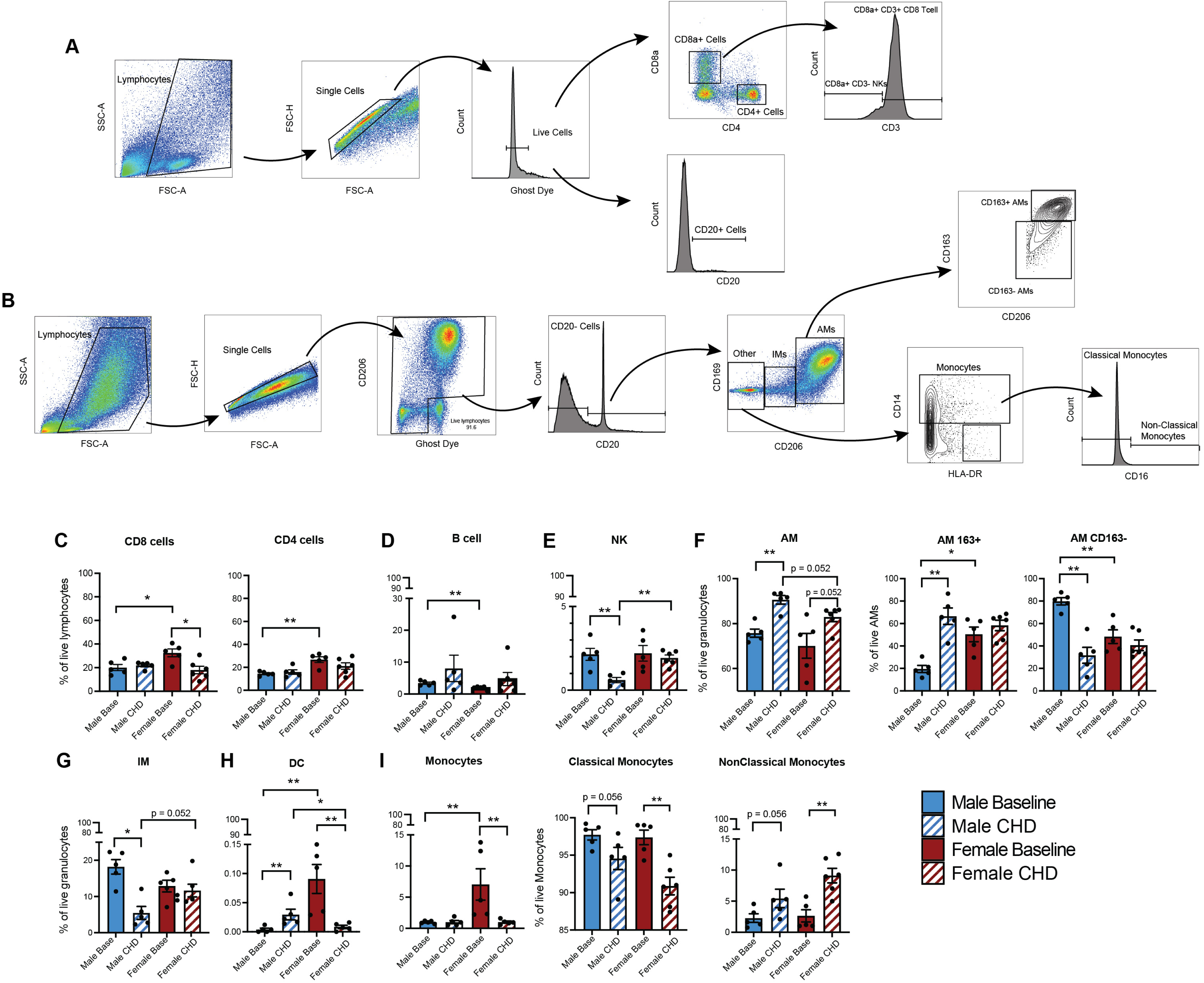
Frequency and gating strategy used to determine immune populations in NHP. Representative flow cytometry gating strategies of **A)** Adaptive and **B)** Innate populations. **C-I)** Barplot of adaptive and innate immune cell population frequencies measured by flow cytometry and stratified by sex and sample timepoint for the indicated populations.

**Supplemental Figure 2:**
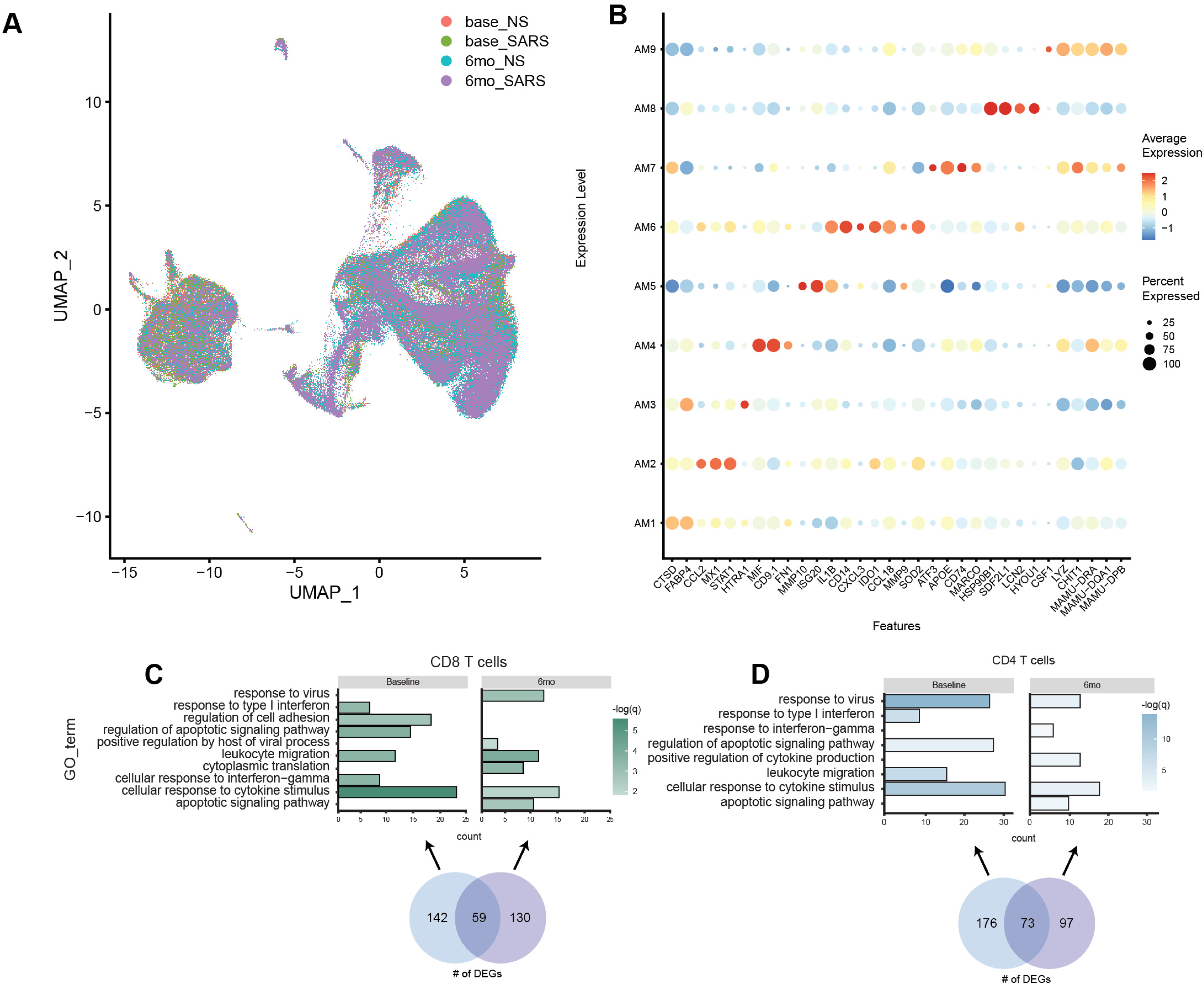
scRNA profiling and DEG analysis of myeloid and lymphoid populations in NHP samples. **A)** UMAP of the immune cells from Figure 1B colored by timepoint and condition. **B)** Bubble plot of marker genes that further differentiate AM clusters from one another. Percent of cells that express each transcript is represented by the size of the bubble while the average expression of those cells is denoted by color, ranging from blue (low) to red (high). **C-D)** Bar graph of notable GO terms from functional enrichment of CD8 (**C**) and CD4 (**D**) T cell clusters. Venn diagrams displaying the number of DEGs unique to and shared between time points are shown as well. Length of the bar is representative of the number of DEGs mapped to corresponding GO term. Color of bar denotes significance.

**Supplemental Figure 3:**
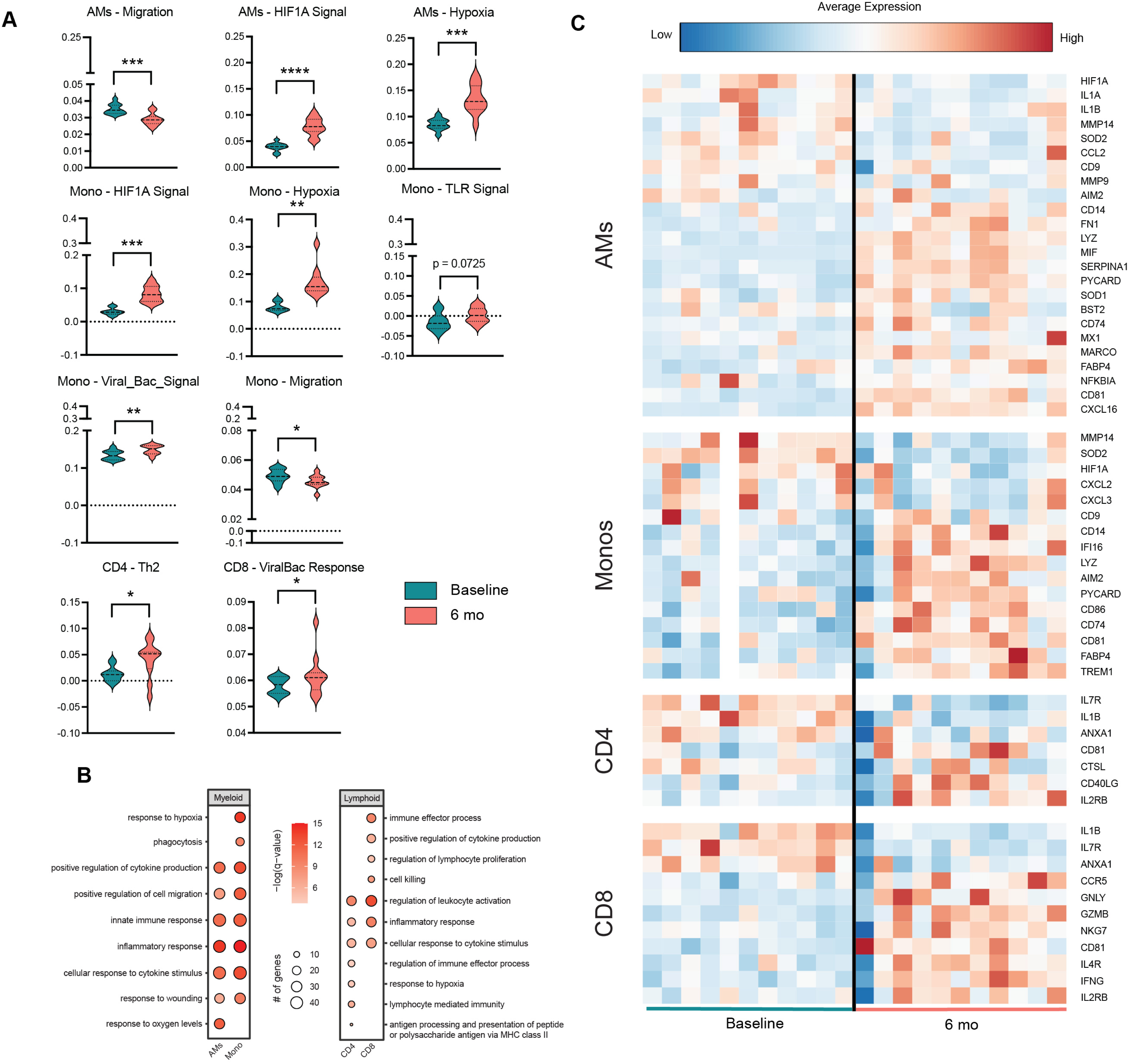
CHD transcriptionally alters myeloid and lymphoid cells in the lung of NHP. **A)** Violin plots comparing module scores of leukocyte activation and chemotaxis (Migration), HIF1A signaling, Hypoxia, TLR signaling, Viral/Bacterial Signaling, and Fc-Receptor mediated phagocytosis for AMs and Mono and Th2 function and Viral/Bacterial Response for CD4 and CD8 T cell clusters respectively at non-infection conditions. Significance calculated using a paired T-test and is denoted as * p ≤ 0.05, **≤ 0.01, *** p ≤ 0.001. **B)** Bubble plot of functional enrichment for select myeloid and lymphoid populations at non-infection conditions. **C)** Heatmap of the average expression of DEGs mapped to GO terms listed in panel B and divided by each monkey in the non-infection conditions at both baseline and EtOH. The average expression of each DEG is denoted by color, ranging from blue (low) to red (high). Note: one of the animals did not contribute to the Mono cluster at baseline.

**Supplemental Figure 4:**
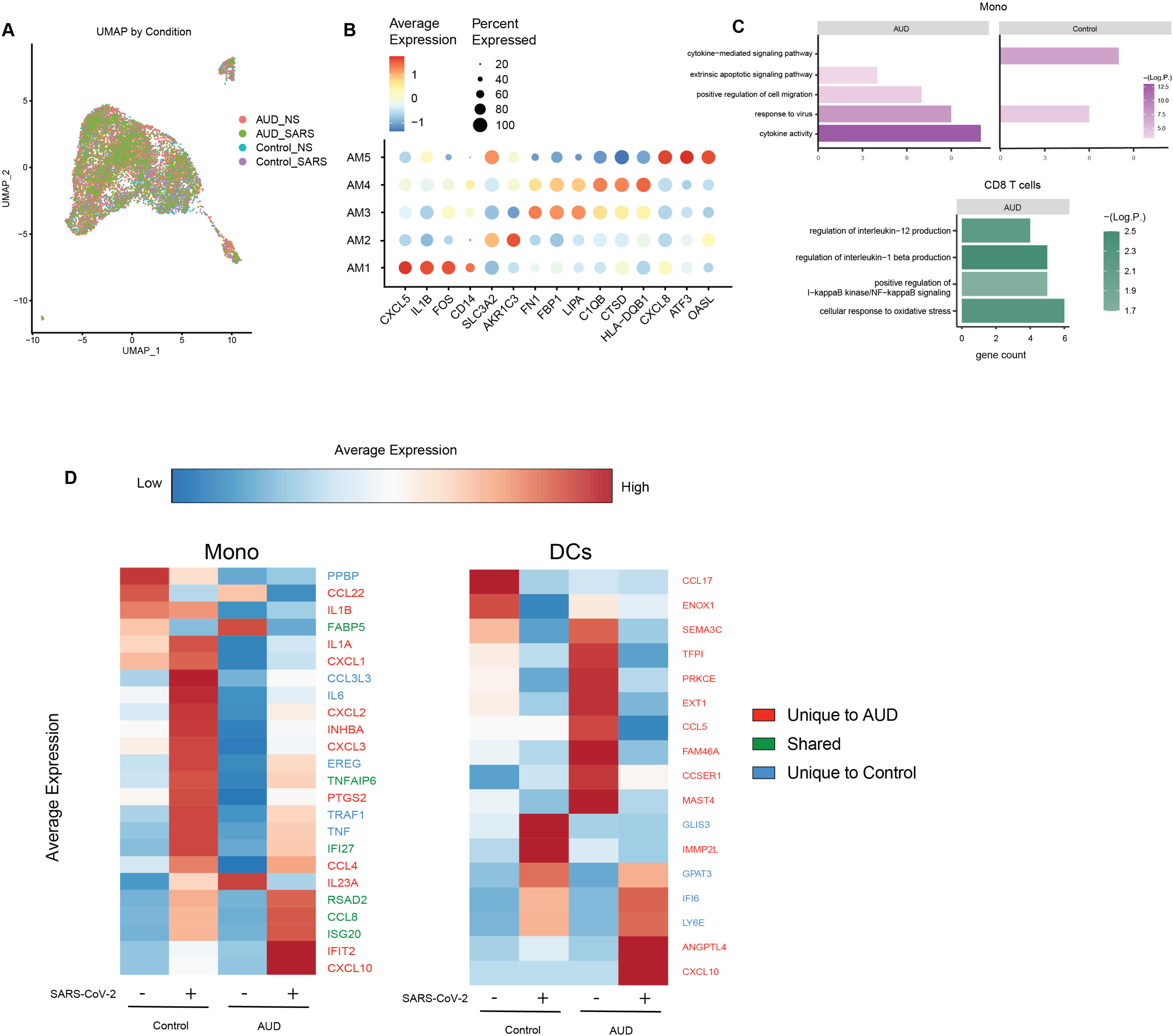
scRNA profiling and DEG analysis of myeloid and lymphoid populations in human samples. **A)** UMAP of cells from Figure 3B colored by condition and timepoint. **B)** Bubble plot of genes that further differentiate the AM clusters from one another. Percent of cells that express each transcript is represented by the size of the bubble while the average expression of those cells is denoted by color, ranging from blue (low) to red (high). **C)** Comparison of GO terms in Mono2 and CD8 T cells, stratified by timepoint. Length of the bar is representative of the number of DEGs mapped to corresponding GO term. Color of bar denotes significance. **D)** Heat maps and violin plots showing gene expression specific to both timepoint and condition. For the heatmaps, the average expression for each transcript ranges from low (blue) to high (red). Genes are colored based on whether they are unique to baseline (blue), AUD (red) or shared between the two (green).

**Supplemental Figure 5:**
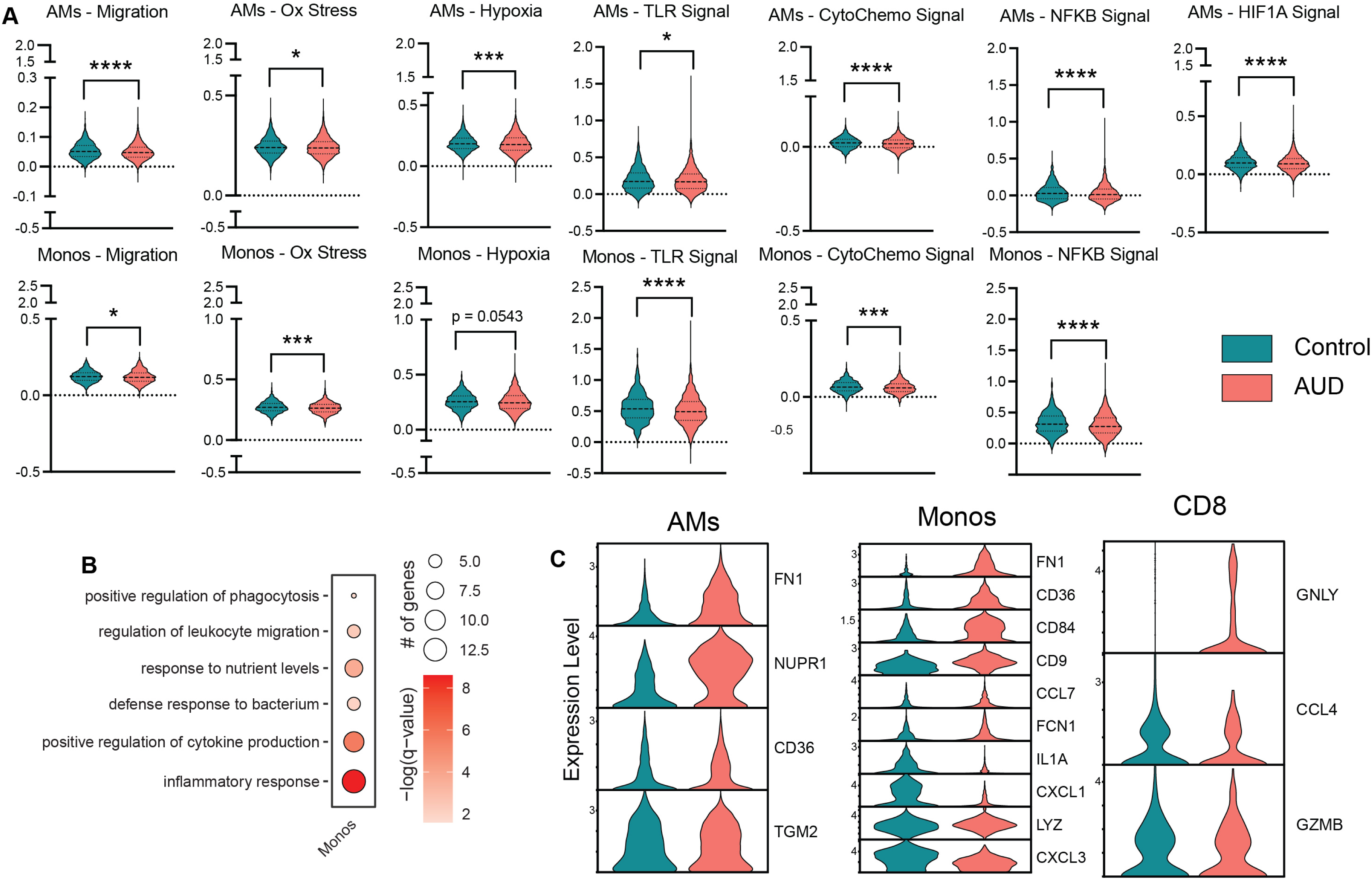
CHD causes transcriptional modifications of immune cells in human samples. **A)** Violin plots comparing module scores of leukocyte activation and chemotaxis (Migration), HIF1A signaling, Hypoxia, Oxyegen stress (Ox Stress), TLR signaling, Viral/Bacterial Signaling, Cytokine/Chemokine Signaling, and NFKB signaling for AMs and monocyte populations (Monos) at non-infection conditions. Significance is denoted as * p ≤ 0.05, ** ≤ 0.01, *** p ≤ 0.001. **B)** Bubble plot of functional enrichment for Monos at non-infection conditions. **C)** Violin plots of the average expression of DEGs for AMs, Monos, and CD8 T cells at both baseline and EtOH. Note: DEGs for Monos were enriched to GO terms listed in panel B. DEGs for AMs and CD8 T cells were not enriched to GO terms.

## Notes

^1^This study was supported by NIH 1R01AA028735-04 (Messaoudi), U01AA013510-20 (Grant), R24AA019431-14 (Grant), and R24AA019661 (Burnham). The content is solely the responsibility of the authors and does not necessarily represent the official views of the NIH.

https://data.mendeley.com/datasets/f44h3v2fn8

https://data.mendeley.com/datasets/729m9vxf4v

### Competing Interest Statement

The authors have declared no competing interest.

