## Supplemental Methods for "Chronic alcohol consumption dysregulates innate immune response to SARS-CoV-2 in the lung"

*Sample collection*

This study leveraged a non-human primate (NHP) model of voluntary ethanol self-administration where macaques had concurrent access to both water and a 4% w/v ethanol solution for 22 hours/day (44). Drinking phenotypes are defined by the pattern and volume of ethanol consumption per day (g/kg/day) and classifed into four categorical levels (LD: low drinking, HD: heavy drinking, VHD: very heavy drinking) defined in (43). Blood samples were collected every 5-7 days, 7 hours into the 22 hours/day drinking session to measure blood ethanol concentration (BEC) via headspace gas chromatography.

For these studies, bronchoalveolar lavage (BAL) samples were collected from 5 male (2 low drinkers and 3 very heavy drinkers) and six female rhesus macaques (4 low drinkers, 1 heavy drinker, and 1 very heavy drinker) at baseline (before induction) and after 6 months of chronic consumption. **Table 1** summarizes the cohort demographics for the rhesus macaques used in this study (Cohort 18 on matrr.com).

Human BAL samples were obtained, with IRB approval, from the University of Colorado School of Medicine’s Colorado-Pulmonary Alcohol Research Collaborative (CoPARC) (<https://medschool.cuanschutz.edu/coparc>) where informed consent was acaquired from all subjects. Three samples were collected from individuals with a history of only smoking and three samples were collected from individuals with a history of chronic alcohol consumption and smoking. **Table 2** summarizes the demographics of the human participants involved in this study.

*Flow Cytometry*

BAL cells were thawed and 1x10^6^were stained with the following antibody panel: CD20, CD3, CD4, CD8a, CD16, CD14, HLA-DR, CD169, CD163, and CD206 (Biolegend) for 30 minutes at 4°C, washed with FACS buffer (2% FBS, 2% 0.5 M EDTA in PBS), and acquired using Attune NxT Flow Cytometer (ThermoFisher Scientific). Results were analyzed using FlowJo software (Ashland, OR).  The remainder of the cells were subjected to SARS-CoV-2 infection.

*Luminex*

Immune mediators of BAL cell culture supernatant were measured before and after stimulation with SARS-CoV-2. NHP samples were analyzed via an R&D 36-plex NHP XL Cytokine Premixed Kit, which contained the following analytes: BDNF, CCL2, CCL5, CCL11, CCL20, CD40 Ligand, CXCL2, CXCL10, CXCL11, FGF basic, G-CSF, GM-CSF, Granzyme B, IFNα, IFNβ, IFNγ, IL-1β, IL-2, IL-4, IL-5, IL-6, IL-7, IL-8, IL-10, IL-12 p70, IL-13, IL-15, IL-17, IL-21, PD-L1, PDGF-AA, PDGF-BB, TGFα, TNFα, VEGF (Bio-Techne, Minneapolis, MN).  Human samples were analyzed via a customized human 29-plex kit containing the following analytes: TNFα, IL-6, PD-L1, PDGF-BB, S100B, IL-7, IFN-β, IL-10, CCL2, VEGF, CXCL13	, IL-1RA, CCL3, CCL4, IL-4, IL-17, IL-2,	IL-15, GM-CSF, IL-8, CXCL9, IFNγ, IL-12P70, IL-1β,	CXCL11, CXCL10, IL-23, CCL11, and IL-18. The samples were analyzed on a MAGPIX instrument (Luminex, Austin, TX). A standard curve was established using the xPONENT™ software and a 5-parameter logistic curve.

*Single Cell RNA library preparation*

After overnight stimulation, BAL cells were washed twice with 0.04% BSA in PBS, before labeling with cell multiplexing oligos (CMOs, 10X genomics) at 4°C for 20 min. Live cells were counted in duplicate and then pooled at a concentration of 1.5x10^6^ cells/mL in 1% BSA in PBS. The pools were stained with Sytox green (Invitrogen) and ~200,000 live cells from each group were sorted using a WOLF cell sorter (Nanocellect Biomedical Inc., San Diego, CA). The sorted cells were resuspended and loaded into a 10x Genomics Chromium Controller at a concentration of 1,600 cells/uL for a target recovery of 20,000 cells. The libraries were prepared using the v3.1 chemistry of the Chromium Single Cell 3’Feature Barcoding Library Kit (10x Genomics, Pleasanton, CA) according to the manufacturer’s instructions.

*Single cells RNA Sequencing Analysis*

The resulting libraries were sequenced using a NovaSeq6000. The reads were aligned and quantified via the Cell Ranger Single-Cell Software Suite and STAR aligner (version 4.0, 10x Genomics) against the Mmul_8 rhesus macaque reference genome for the macaque samples and the human reference genome GRCh38 for the human samples. Downstream analysis was performed separately using Seurat (version 4.1.1). Briefly, samples were de-multiplexed based on their unique CMOs followed by the removal of ambient RNA (<200 feature counts) and dead cells (>5% mitochondrial gene expression). The samples were normalized and scaled based on cell cycle and mitochondrial gene expression using *NormalizeData* and *ScaleData,* respectively. Data sets were integrated using reciprocal PCA via Seurat’s *FindIntegrationAnchors*. Once the anchor points were found, the samples were combined into a single data frame using *IntegrateData*. These integrated data were again scaled using *ScaleData,* and a PCA was generated to determine the dimensionality of the data set via *RunPCA*. UMAP generation and cell clustering was performed using *RunUMAP* (dims = 1:50), *FindNeighbors* (dims = 1:20), and *FindClusters* (resolution = 0.5). The identity of the resulting clusters was then determined using the canonical markers found using *FindAllMarkers* and a Log2 fold change cutoff of 0.4. Differential gene expression analysis was performed using the default settings of MAST in Seurat and comparing SARS-CoV-2 infected to non-infected cells at each timepoint. Differentially expressed genes (DEGs) were defined as those with FDR p ≤ 0.05 and Log2 fold change ≥ 0.25. When comparing infection to non-infection in human samples, the Log2 fold change cut-off was removed. Functional enrichment was performed through Metascape.

*Statistical Analysis*

Statistical analyses for non scRNA-Seq data sets were performed using the GraphPad Prism software (GraphPad Software Inc., La Jolla, CA). All analyses were tested for Gaussian distribution using a Shapiro-Wilk normality test (α = 0.05). Subsequent tests were performed non-parametrically the data set failed normality. If possible, statistical analyses were performed in a pairwise fashion. When comparing more than two groups, we used a 1-way analysis of variance (ANOVA) with Holm-Šídák *post hoc* multiple comparisons or Friedman test with Dunn’s *post hoc* multiple comparisons. For two groups, non-parametric T tests were carried out using Mann-Whitney test or Wilcoxon matched-pairs signed ranked test.
